# Disentangling the impact of environmental and phylogenetic constraints on prokaryotic strain diversity

**DOI:** 10.1101/735696

**Authors:** Oleksandr M. Maistrenko, Daniel R. Mende, Mechthild Luetge, Falk Hildebrand, Thomas S. B. Schmidt, Simone S. Li, Luis Pedro Coelho, Jaime Huerta-Cepas, Shinichi Sunagawa, Peer Bork

## Abstract

Microbial organisms inhabit virtually all environments and encompass a vast biological diversity. The pan-genome concept aims to facilitate an understanding of diversity within defined phylogenetic groups. Hence, pan-genomes are increasingly used to characterize the strain diversity of prokaryotic species. To understand the interdependency of pan-genome features (such as numbers of core and accessory genes) and to study the impact of environmental and phylogenetic constraints on the evolution of conspecific strains, we computed pan-genomes for 155 phylogenetically diverse species using 7000 high-quality genomes. We show that many pan-genome features such as functional diversity and core genome nucleotide diversity are correlated to each other. Further, habitat flexibility as approximated by species ubiquity is associated with several pan-genome features, particularly core genome size. In general, environment had a stronger impact on pan-genome features than phylogenetic signal. Similar environmental preferences led to convergent evolution of pan-genomic features in distant phylogenetic clades. For example, the soil environment promotes expansion of pan-genome size, while host-associated habitats lead to its reduction.

## Introduction

Sequenced prokaryotic species vary approximately 100 fold in genome size and gene content [1]. The gene content of bacterial and archaeal genomes is mainly shaped by gene duplication, neo-/sub-functionalization, and losses. Other sources of functional innovation include *de novo* emergence of genes, and horizontal transfers; all leading to a vast prokaryotic genetic diversity [2–4]. In order to characterize strain diversity within a species, pan-genome analyses have been proven useful [5]. The pan-genome is the non-redundant set of all genes (gene clusters or homologous groups) found in all genomes of a taxon [6, 7]. A species pangenome contains core genes (that are present in almost all isolates) and accessory genes, which can be further subdivided based on their prevalence. Each newly sequenced genome of a conspecific strain can contribute anywhere between 0 to more than 300 new genes to the pan-genome of a species [8]. This potentially infinite addition of new genes means that the accessory gene repertoire of a species can theoretically increase with no emerging upper boundary, making pan-genomes appear *open*.

The pan-genome of a given species is potentially shaped by its respective habitat(s) (via selection and drift) and phylogeny (inherited gene content after speciation). For example, previous studies have observed a relationship between habitat and genome size (as a proxy for gene content): free-living soil bacteria tend to have the largest described genomes [9] while marine free-living and intracellular symbionts harbor the smallest ones [10–13]. Obligate symbiotic species tend to have small pan-genomes – almost equal to the genome size, while soil-associated and some highly abundant free-living marine bacteria tend to have the largest pan-genomes [14]. However, it is not well understood which aspects of a species’ pangenome are influenced by environmental factors and phylogenetic inertia. The overall architecture of a pan-genome can be described from various angles, using established quantitative measures of individual pan-genome features, such as pan-/core genome genome sizes, genome fluidity, and average nucleotide identity/diversity (also see Supplementary Table 1 for definitions). Many of the pan-genome features describe the size of certain categories of genes, while others focus on a description of within species diversity.

Pan-genome features are generally expected to be phylogenetically conserved as a result of the evolutionary history of a given species, and predefined by past exposures to different environments. Closely related species tend to share more genes, i.e. gene content similarity follows phylogeny [2, 15]. Further, habitat preferences are also phylogenetically predetermined [16] and dispersal capability varies across different taxa [17, 18]. On the other hand, environmental factors shape genome architecture and the pan-genome in general [19]. A (pan-)genome’s functional potential mirrors both niche and phylogenetic signals [20], consequently phylogenetic relatedness and genome functionality are mildly predictive of species ubiquity and genome size [21, 22]. Thus, pan-genome features exhibit both phylogenetic and environmental associations. Yet as phylogeny and habitat preference are themselves correlated, their association needs to be taken into account when disentangling their relative contributions to pan-genome features (Fig. 1).

**Fig. 1.**
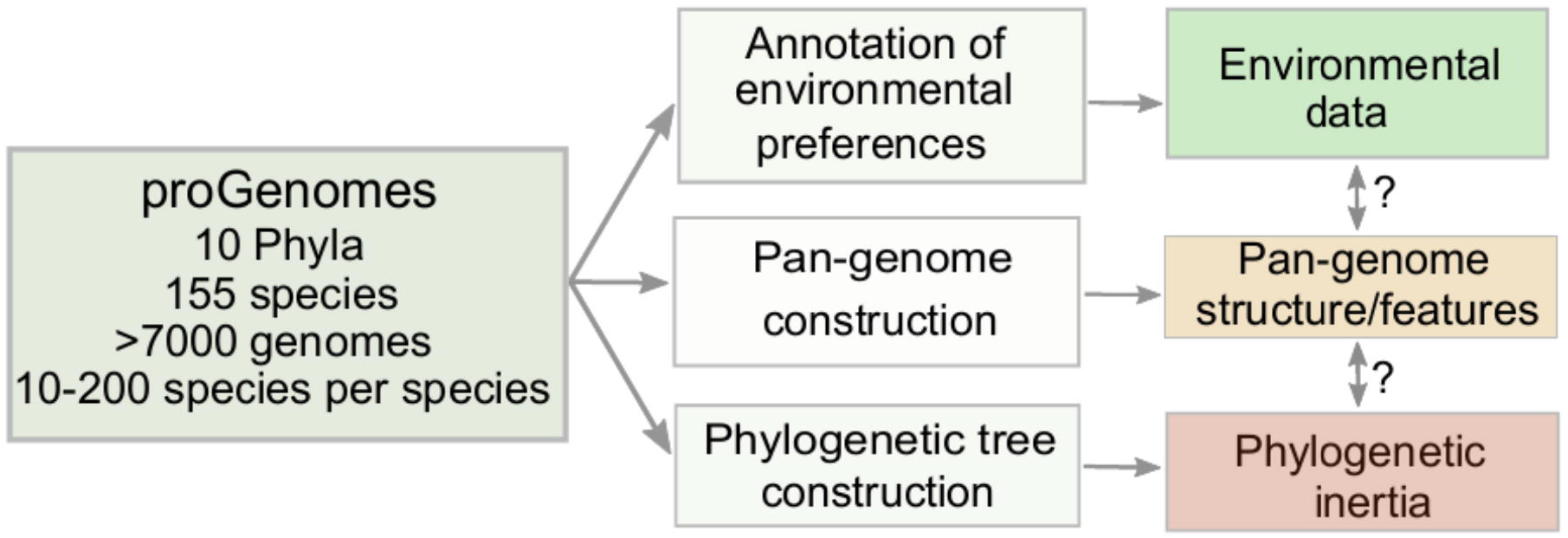
Study design. We used the proGenomes database v.1 [28] of high-quality genomes to compute pan-genomes, pan-genomic features and assigned preferred habitats to species (see methods). As many pan-genome features are interdependent (covariates) or affected by sampling bias, we used a multivariate analysis framework to disentangle habitat properties from phylogenetic inertia. This allows for the quantification of environmental and phylogenetic factors that impact strain diversity within species.

Recently, the pan-genome and its derivative measures (features) have been used extensively in comparative genomics of individual prokaryotic species to: (i) define species boundaries [23, 24], (ii) describe the genomic diversity of species [25], (iii) reveal origins of mutualistic and pathogenic strains [14] and (iv) characterize evolutionary and ecological mechanisms that shape genome architecture [8, 26]. To understand the general principles of pan-genome evolution and to disentangle the impact of environment and phylogeny, we performed a meta-analysis of over 7000 genomes, encompassing 155 prokaryotic species from 10 phyla and 83 environments (Fig. 1). We computed 21 previously established pan-genomic features. The variation across these features was explored with respect to phylogenetic inertia and environmental constraints/preferences (as characterized by 83 habitat descriptors) of the studied species. Using this framework, we quantified interdependencies of pan-genomic features, identified novel relationships among them, and estimated how environment shapes pan-genome architecture.

## Methods

### Genomic data

In this study we used 7104 genomes from 155 species (defined using 40 universal marker genes - specI clusters [27]) obtained from the proGenomes database [28] (see Supplementary Table 2). For reliability of further analysis, we included only high-quality genomes with 300 or fewer contigs. Only one genome from any pair of genomes was retained for downstream analysis when pairwise nucleotide identity in core genome was 100% and pairwise gene content overlap (Jaccard index) > 99%. We used only species that contained at least 10 high-quality genomes in the proGenomes database [28].

### Habitat annotation

Habitat metadata for isolates/strains were obtained from the PATRIC database [29], the Microbe Atlas Project database (JF Matias Rodrigues et al, *in preparation*) and Global Microbial Gene Catalog (LP Coelho et al, *submitted*), resulting in the reliable annotation of species to one or more habitats (83 total habitats). Ubiquity was estimated as the sum of all positive associations (Fisher’s Exact Tests, Benjamini-Hochberg-correction, p≤0.05) across all habitats in the Microbe Atlas Project dataset. The final annotation is available as Supplementary Table 3.

### Pan-genome reconstruction

Pan-genomes for the 155 species studied were constructed using the Roary pipeline [30]. Input genomes for pan-genome construction were first annotated using Prokka [31]. We identified homologous gene clusters at an amino acid identity threshold of 80% [32–35]. Pan- and core genome curves were generated via 30 input order permutations (similar to the approach in the GET_HOMOLOGUES pan-genome pipeline [36]. Fitting of non-linear regressions was performed in R v.3.3.2 [37] using “nls package”. The total number of genes in the pan-genome of a given species, the number of new genes added per genome and the total number of core genes were modeled using equations [1], [2] and [3] respectively to estimate openness of pan-genome [6, 7].

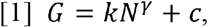

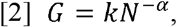

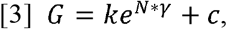

where G – number of genes; N – genome number that is added to analysis; k, c, - constants; α and γ – saturation coefficients. When γ≤0 in equation [1] – pan-genome is closed (saturated) (Fig. 2a); 0≪γ≤1 – pan-genome is open (Fig. 2a). When α<1 in [2] – pan-genome open, α>1 – pan-genome is closed.

**Fig. 2.**
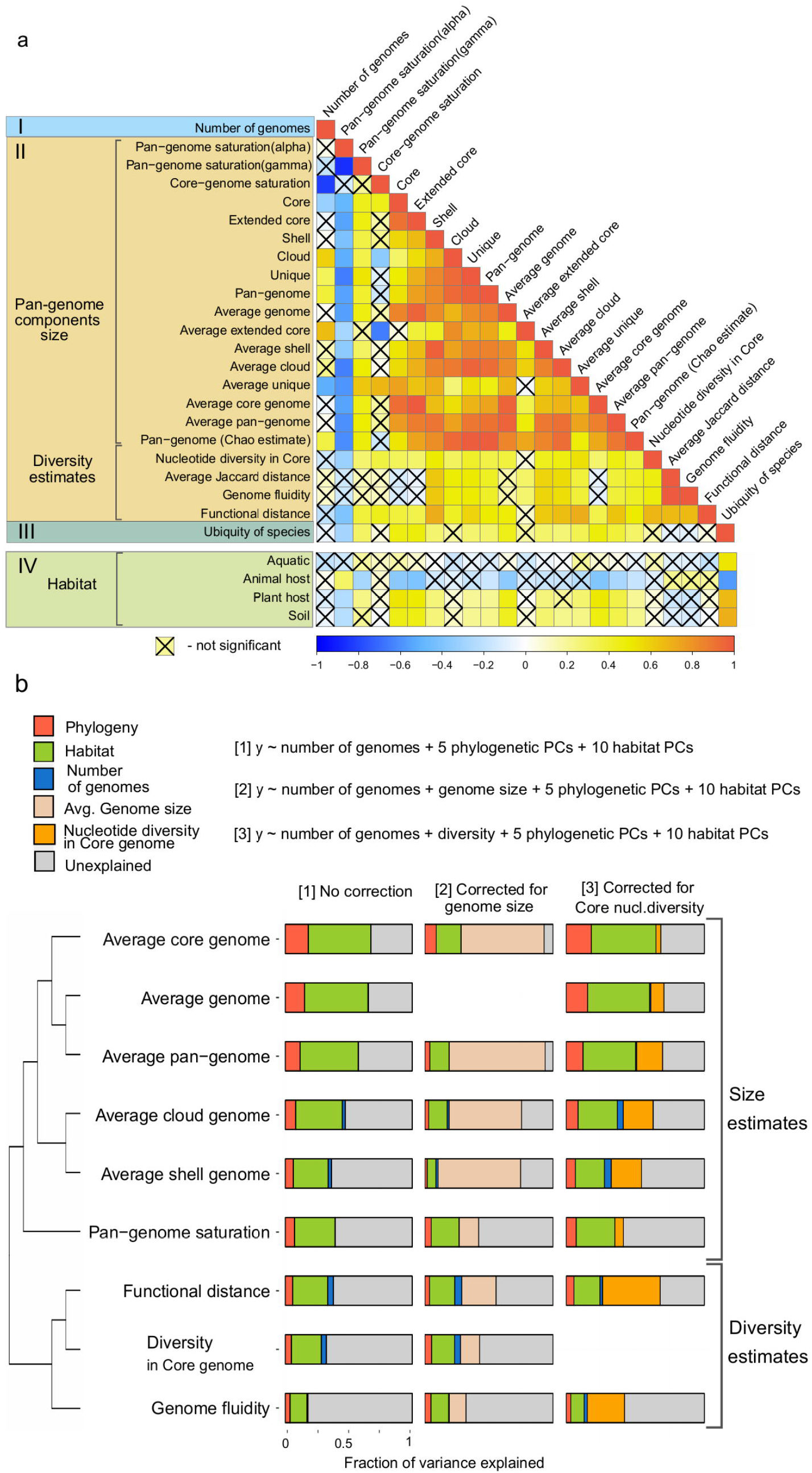
Relationship between different pan-genome features. **a**. Correlation matrix between (I) the number of conspecific genomes used to estimate pan-genome features, (II) 21 pan-genome features, (III) the ubiquity of species as an environmental feature computed from habitat preference of strains, and (IV) major habitat groups from the Microbial Atlas project. The heatmap visualizes Spearman Rho values for correlations between sample size (I), 21 pangenome features (II) and species ubiquity (III). Four major habitats (aquatic, animal host, plant host, soil (IV)) were correlated to the (I) number of conspecific genomes, (II) pangenome features and (III) ubiquity via point-biserial correlation. Statistical significance of correlations was determined using adjusted p-values (using Benjamin-Hochberg correction) < 0.05. **b**. Clustering of a subset of nine pan-genome features based on their pairwise correlation strengths. Horizontal stacked charts present amount of variance explained by various predictors (number of genomes, phylogeny and habitat represented by their principal components (PCs), and genome size or diversity). The first set of stacked charts (“no correction”) shows variance explained in pan-genome features by the number of genomes used to compute pan-genome features as well as species’ phylogeny and habitat preferences; the second and the third sets of stacked charts represent the amount of variance explained (see Methods) by the same set of predictors when correcting for genome size or nucleotide diversity in core genome respectively. Size and diversity estimates form distinct feature groups.

Classification thresholds for pan-genome subcomponents were defined as follows: core genes – present in all strains; extended core – present in > 90% of genomes; cloud genes – present in < 15% (includes unique genes in pan-genome); the remaining part of pan-genome were considered “shell” genes (Supplementary Figure 1). These thresholds are based on default parameters of the Roary pipeline [30], although we readjusted the extended core threshold to 90%, as suggested by the distribution frequency of genes within the pan-genomes in our dataset (Supplementary Figure 2). The R package “micropan” [38] was used to compute genomic fluidity [39], Chao’s lower bound for gene content in the pan-genome [40] and Heaps’ alpha (equation [2]) [6]. Functional distance between strains within each pangenome was estimated as jaccard distance based on eggNog v4.5 annotations [41] of pangenome gene clusters. 23 parameters (21 pan-genome features, plus the number of conspecific isolates and species ubiquity) were compared using Spearman’s rank correlation to investigate the relationship between sample sizes, subcomponents of pan-genome, saturation parameters (γ and α) from equations [1], [2], [3], genome fluidity functional distance and core genome nucleotide identity (see Supplementary Table 1 for definitions of pan-genome features). To obtain unbiased estimates of core and pan-genome size we rarefied to 9 genomes as suggested previously [36]. Hierarchical clustering of a subset of pan-genome features was performed on absolute values of pairwise Spearman Rho values as displayed in Fig. 2a.

### Phylogenetic signal and phylogenetic generalized least squares

An approximate maximum likelihood phylogenetic tree of all 155 species was generated using the *ete-build* concatenation workflow “clustalo_default-trimal01-none-none” and “sptree-fasttree-all” from ETE Toolkit v3.1.1 [42], using protein sequences of 40 conserved universal marker genes [27, 43, 44] and default parameters for the ClustalOmega aligner [45] and FastTree2 [46] with the JTT model [47].

To estimate the phylogenetic signal of genomic traits, we used the R package “phylosignal” [48] with Pagel’s Lambda [49], following guidelines for phylogenetic signal analysis [50, 51] (Supplementary Figure 3). The “Caper” R-package was used for phylogenetic generalized least squares regression [52].

### Variance quantification

The cophenetic distance matrix obtained from the phylogenetic tree and the binary habitat association matrix were each decomposed using the “FactoMineR” R package [53]. The first 5 phylogenetic principal components (PCs, accounting for ∼80% of phylogenetic variance) and 10 habitat PCs (accounting for ∼50% of habitat variance) were used for variance partitioning. PCs were selected using the “broken stick” model utilizing a computer program made available by Borcard et al 2011 [54]. The first two principal components for phylogenetic and habitat matrices decompositions are visualized in Supplementary Figure 4 and Supplementary Figure 5. The fraction of the variance explained by habitat and phylogeny were estimated using the CAR metric which performs a decorrelation of predictors [55] implemented in the “car” R-package with the following models:

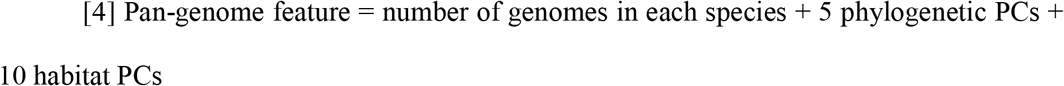

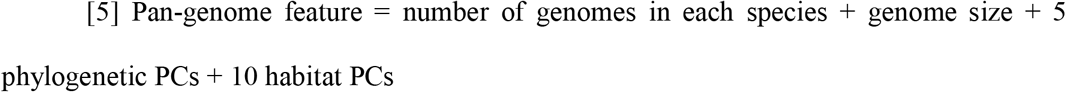

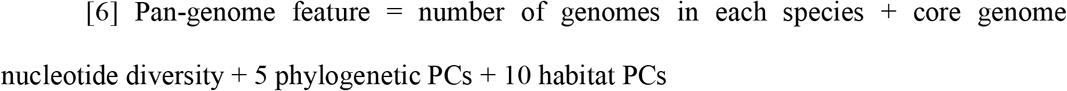

We also performed the model fitting procedure [4] on 1000 permutations of the first 5 phylogenetic PCs and first 10 habitat PCs to ensure that the actual habitats and phylogeny data explained a higher fraction of the variance than randomized models (Supplementary Figure 6).

## Results

### Delineation of pan-genomes and habitats descriptors

The foundation of this study is a large collection of pan-genomes from a diverse set of prokaryotic species. To establish this foundation, we filtered the proGenomes database of annotated prokaryotic genomes [28] to select species (see Methods, also [27]) for which at least 10 high-quality genomes (conspecific isolates/strains/genomes; further referenced as strains or genomes) were available (Fig. 1, also see Methods). For each of the resulting 155 species, we computed 21 pan-genome features (ranging from pan-genome saturation to functional distance, see Fig. 2a and Supplementary Table 1). These features have been shown to characterize different aspects of the pan-genomic structure and have been previously used in pan-genome analysis projects on individual microbial species [6, 39]. Partitioning of the pan-genome into subcomponents (“core”, “shell”, “cloud”; see Methods) enabled us to relate the evolutionary adaptations of core and accessory genome features to environmental pressures separately. Pan-genome features varied in size, for example, average core genome size was in the range of 443-5964 genes; average pan-genome size – 959-17739 genes; average shell 18-2409; average cloud – 5-839 genes. We annotated each species’ habitat preferences (operationally defined as the set of habitats where species were observed) by merging information obtained from the PATRIC database [29], the Microbial Atlas Project database (http://devel.microbeatlas.org/ see also Supplementary Table 3) (JF Matias Rodrigues et al, *in preparation*) and the Global Microbial Gene Catalog (https://gmgc.embl.de) (LP Coelho et al, *submitted*), resulting in assignments of each genome (strain) to one or multiple of 83 habitat descriptors (see methods). On average, each species was present in 16.5±7.8 habitats in Microbial Atlas Project (JF Matias Rodrigues et al, *in preparation);* 2.4±1.1 from manually curated PATRIC habitat annotations; and 3.6±2.8 in the Global Microbial Gene Catalog (LP Coelho et al, *submitted*). (Supplementary Table 3).

### Interdependencies of pan-genome features

The relationships between different pan-genomes features can be an indication of similar evolutionary pressures acting on the related features. Further, correlations between different features can decrease the accuracy of analyses when not considered. Hence, we estimated interdependencies for (i) the number of conspecific strains (the number of genomes per pan-genome), (ii) the 21 computed pan-genome features, (iii) species ubiquity, and (iv) habitat preference (see Supplementary Table 3 for estimates of pan-genome features, Supplementary Table 1 for definition and Supplementary Table 4 for correlation summary) (Fig. 2a). Estimates of pan-genome size and the size of its components (core, shell, cloud) are strongly correlated with each other (Fig. 2a). As expected, mean genome size strongly correlated with several features, including core-genome size (Spearman Rho 0.955, p<0.00001), pan-genome size (0.963, p<0.00001) and core-genome nucleotide diversity (0.373, p=0.00003), indicating that a species’ genome size is highly predictive of its pangenome features, especially pan-genome size.

Unexpectedly, several pairs of conceptually related pan-genome features did not correlate. For example, the correlation between genome fluidity (average ratio of unique gene clusters/families to the sum of gene families over random subset pairs of genomes, see Supplementary Table 1 [39] and pan-genome saturation (a representation of the number of new genes are expected for every new genome sequenced, see Methods) was not significant (0.15, p=0.72), despite the fact that both measures are commonly used to estimate the openness of pan-genomes [8, 39]. This might indicate that these two measures characterize different aspects of pan-genome openness. A possible explanation of this observation is an implicit bias of the pan-genome saturation estimate due to under-sampling [39].

Furthermore, the average pairwise functional distance (average Jaccard distance based on orthologous groups) between conspecific strains positively correlated to the vast majority of pan-genome features. Only three pan-genome features were not significantly correlated to the average pairwise functional distance, namely the size of the extended core, the number of conspecific strains (number of conspecific genomes used to compute pan-genome features) and ubiquity (see Supplementary Table 4 for Spearman Rho and p-values). We also found that species with larger genomes tend to have a higher functional diversity (Spearman Rho = 0.48, *p* = 6.5 * 10^−9^), mainly driven by changes in the size of the pan-genome shell. This relationship is not biased by the number of sampled strains (Fig. 2a).

We observed associations of sample size (number of conspecific strains) with core genome saturation, total pan-genome size, and the sizes of “cloud”, “unique genes”, and “extended core”, indicating a sampling bias for a subset of the species analyzed. To compensate for this potential bias, we performed variance partitioning on 9 out of 21 features representing qualitatively different pan-genome properties are not significantly affected by sample size (non-significant correlations with Spearman Rho close to 0, see Fig. 2a). We explored the interdependencies of these nine pan-genome features by clustering them according to their correlation strengths and identified two subgroups (Fig. 2b, see also Supplementary Table 5). These subgroups split the features into diversity estimates (core genome nucleotide diversity, functional distance, genome fluidity) and size estimates (average genome, pan-genome, core, shell, cloud), implying differing evolutionary dynamics for these feature groups. Specifically, size-related pan-genome features were better explained by phylogenetic and environmental preference compared to diversity estimates (Fig. 2b).

### Species ubiquity is related to core genome size

All surveyed species are abundant in multiple habitats and the transition between free-living and host-associated lifestyles is frequent on both micro- and macroevolutionary (and ecological and evolutionary) timescales, imposing multidirectional pressures on the evolution of their genome architecture [56]. Species ubiquity is a potentially important factor contributing to the evolution of specific pan-genome features that needs to be considered, because species with broad ecological niche are likely to have different evolutionary constraints compared to specialists [57]. We operationally defined species ubiquity as the sum of all positive associations with each habitat in the Microbe Atlas Project dataset (see Methods). Broader ecological niches and higher ubiquity tend to be associated with larger and more functionally versatile genomes [58]. Therefore, we investigated the relationship between the ubiquity of each species with its pan-genome features in depth and found several associations (Fig. 2a). We observed a moderate, but significant association of species ubiquity (Fig. 3a) with average normalized core genome size (operationally rarefied to 9 genomes to obtain an unbiased average of core genome size and its s. d., see Supplementary Figure 1) and on pan-genome saturation (exponent coefficient α in Heap’s law model, equation [2] in Methods), but not on any other pan-genome features after correcting for phylogenetic effects (Fig. 3a, Supplementary Table 6). This suggests that a larger core genome may be important to facilitate persistence and proliferation in multiple habitats. The core genome of highly ubiquitous species was enriched in genes coding for proteins involved in lipid metabolism and secondary metabolite biosynthesis (I and Q in Fig. 3b, respectively). This is congruent with earlier studies, suggesting that secondary metabolite biosynthesis might be implicated in adaptation to multiple environments [58].

**Fig. 3.**
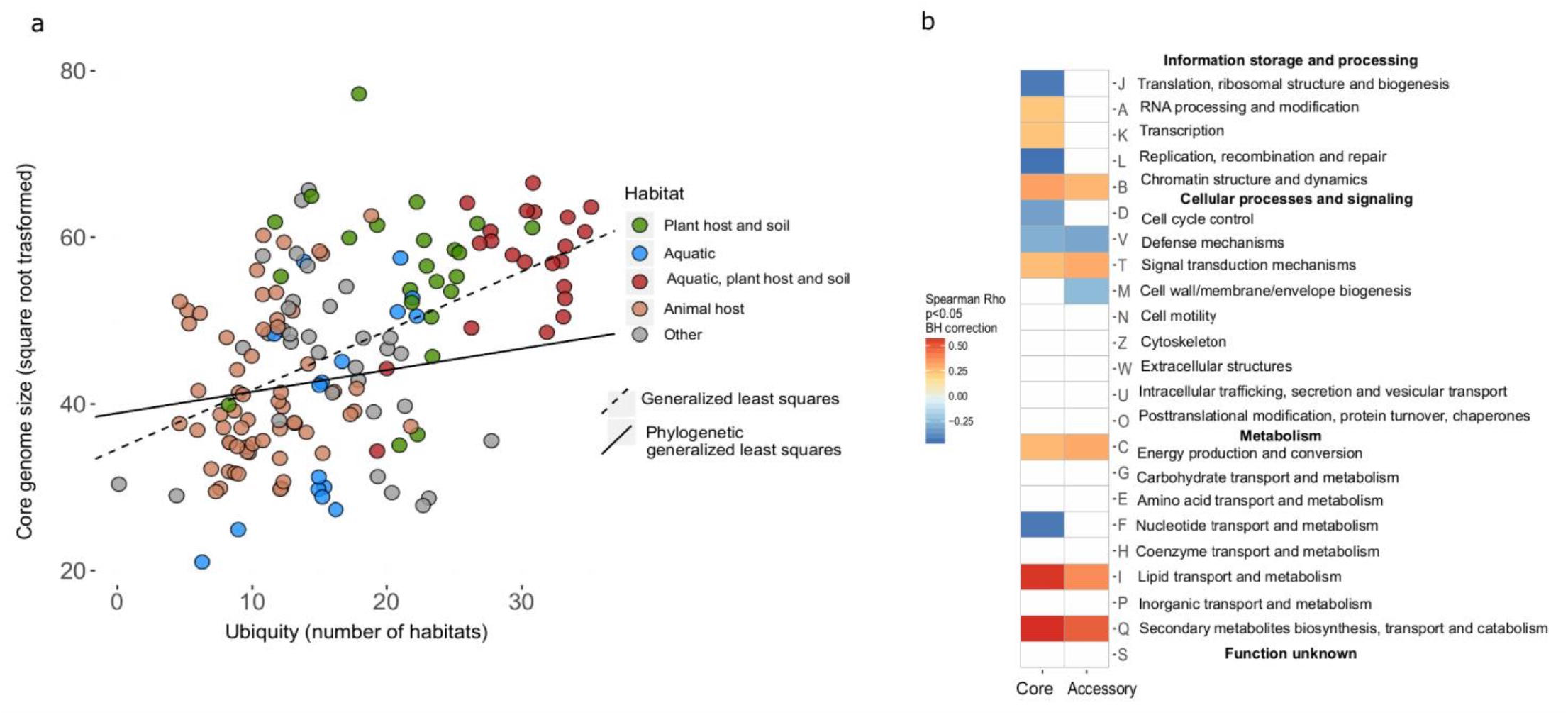
Effect of ubiquity on core genome size and functional content. **a**. Species ubiquity (number of habitats a species was assigned to), a habitat feature, is linked to core genome sizes after correction for phylogenetic effect (Phylogenetic generalized least squares, p-value=0.00005, λ=0.98 (95% C.I. 0.957, 0.992), partial r^2 (for ubiquity coefficient) 0.09, see also Supplementary Table 6). **b**. Correlation of ubiquity with the relative frequency of functional categories (COG categories assigned by eggNog v4.5 [41]) in core and accessory genomes. Species of high ubiquity tend to encode more proteins involved in Lipid Metabolism (I) and Secondary Metabolite Biosynthesis (Q).

### Dissecting the impact of phylogenetic inertia and environment on pan-genome features

We next quantified differential contributions of evolution and environmental factors to pan-genome architecture. Pan-genome features were modeled as a combination of the number of conspecific genomes considered, phylogenetic placement and habitat preference. For this we used an abstract representation of phylogeny and habitats as principal components (PCs), accounting for dimensionality, collinearity and redundancy within these data. The respective relationships were approximated using a linear model (Methods), which allowed us to estimate the variance of pan-genome features between species explained by phylogenetic effect and habitat preferences:

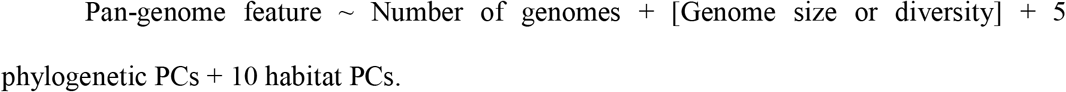

Together, habitat and phylogenetic effects explained large parts of the variance (up to 49% by habitat and 17% by phylogenetic effect) in all selected features (Fig. 2b, Supplementary Table 5). This remained true, even when controlling for genome size or diversity of the core genome (as evident when these were included in the model as predictors as in the second and third set of stacked charts of Fig. 2b) (Supplementary Table 6). Habitat and phylogeny have considerable independent effects on pan-genomic features, although the impact of habitat preferences was consistently stronger (Fig. 4). Diversity estimates, in contrast, were explained to a lesser degree by habitat preferences of species and phylogenetic inertia, as they likely reflect spatio-temporal (microevolutionary) variation of sub-populations within species due to local adaptation and/or genetic drift [25, 59]. For example, a higher fraction of core genome size (and genome size) variance was explained by species habitat preference than any other pan-genome feature (including accessory genome size when considered separately), implying that core genome size might be linked to a species’ ecology while the accessory genome might often be more affected by random gene acquisition and loss. The observed signals were robust to technical and annotation noise, as random permutations of habitat and phylogenetic principal components did not exceed the observed data in variance explained (except for genome fluidity (Supplementary Figure 6)). The strongest phylogenetic effects were observed for average core, pan-genome and genome sizes (confirmed using Pagel’s Lambda estimate to test the strength of the phylogenetic signal [49] (Supplementary Figure 3). Altogether, up to 65% of the variance of different pan-genome features was explained by habitat and phylogeny (Fig. 2b and Fig. 4). Overall, habitat and phylogeny effects contributed differentially to diversity- and size-based pan-genome features (Fig. 2b).

**Fig. 4.**
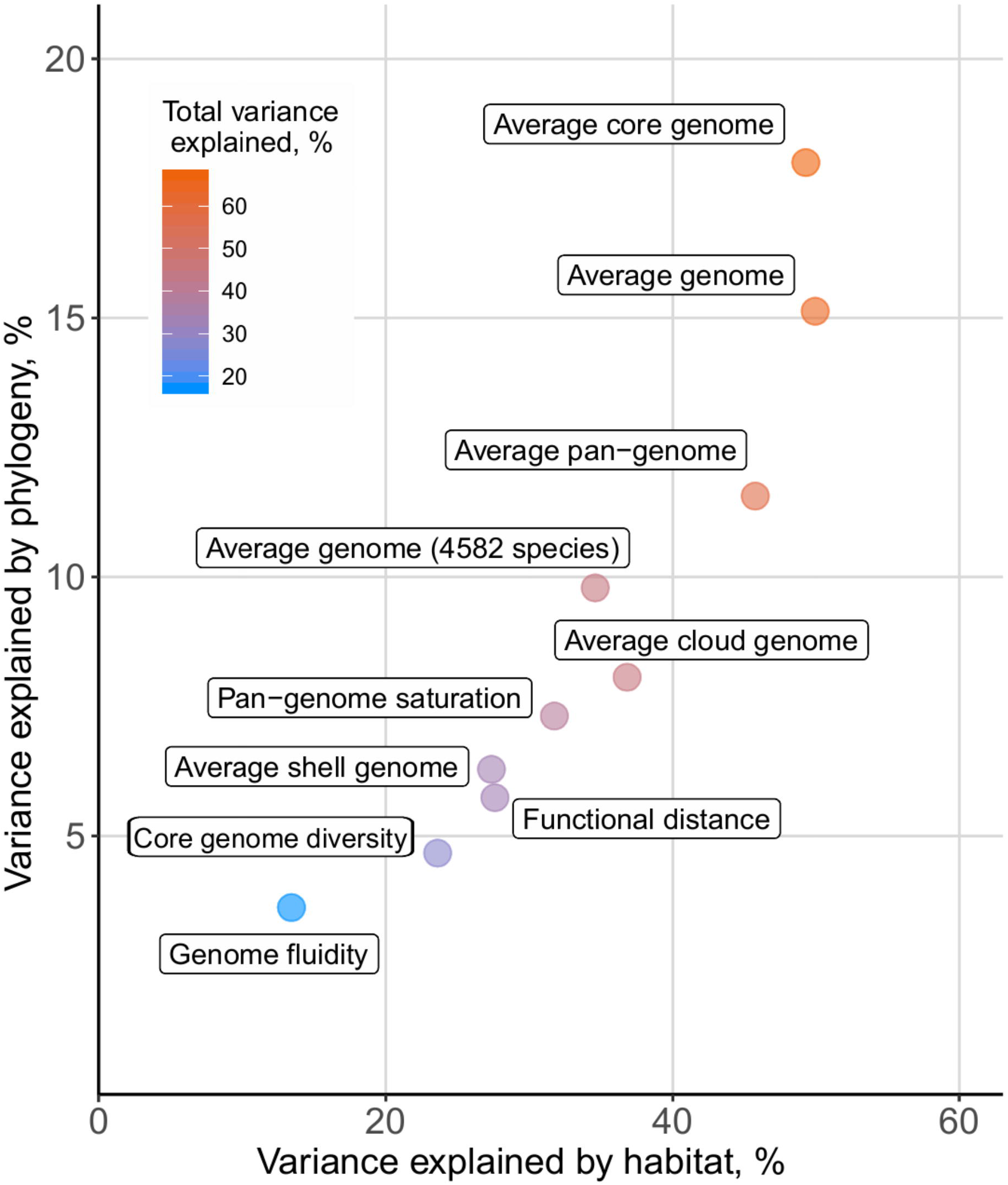
Partitioning of variance in pan-genome features by phylogeny and habitat (R-square(car score)) based on model [1] from Fig. 2b.

### Environment-driven, convergent evolution of pan-genome features

We next tried to narrow down the habitat effects of selected major habitat groups (soil-associated, aquatic, animal-host associated and plant-host associated habitats) in relation to genome/pan-genomes sizes and diversity, accounting for phylogenetic background (Fig. 2a). Soil and plant-host habitats were associated with larger pan- and core genomes, while animal host habitats were associated with smaller ones. Aquatic habitats were not a good predictor for size-related pan-genome features, which might be indicative of their heterogeneous nature [19, 60]. The distribution of core genome sizes across the phylogenetic tree of species studied showed that large core genomes have independently evolved (Kruskal-Wallis test, chi-squared = 32.194, df = 1, p-value = 1.395e-08) in soil-inhabiting species from at least four (out of ten analyzed) phyla (Proteobacteria, Actinobacteria, Spirochaetes and Firmicutes, Fig. 5). Small core genome sizes independently evolved at least 3 times (Proteobacteria, Actinobacteria, Firmicutes) in our dataset. Nucleotide diversity of the core genome was, in contrast to size, less affected by habitat and phylogenetic signals (Fig. 4, Supplementary Figure 3). Nevertheless, species with a higher nucleotide diversity within their core genome were positively associated with aquatic habitats (Fig. 5) (Kruskal-Wallis test, chi-squared = 25.69, df = 1, p-value = 4.01e-07), in line with earlier observations from metagenomics [16]. In conclusion, core genome sizes and (to a lesser degree) diversity in prokaryotic species depend on broad habitat type(s) and range, implying that adaptation to a given habitat range leads to convergent evolution towards habitat-specific core genome sizes (e.g. soil-associated species have larger genomes, Fig. 5). Our findings were consistent when using larger cohort of over 4500 species for which we explored the relationships between genome size and habitat as well as phylogeny (Fig. 4, also see Supplementary Table 6 and Supplementary Table 7). As expected, habitat had a much greater effect (34.6% variance explained) than phylogeny (9.8% variance explained). This implies that our findings across 155 diverse species (Fig. 3a) are likely to be generalizable.

**Fig. 5.**
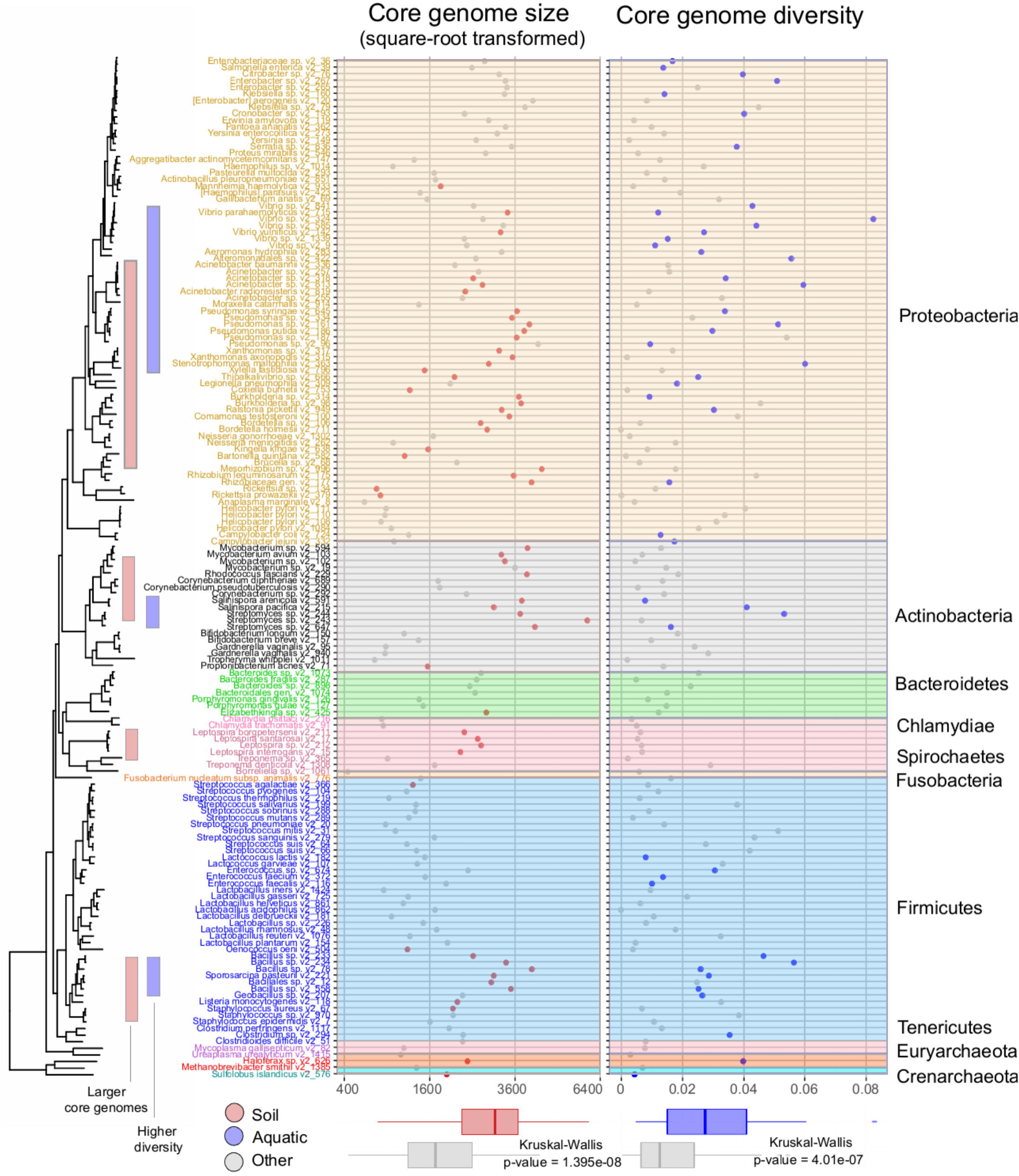
Phylogenetic tree of 155 microbial species with scatterplots of core genome size and average nucleotide diversity of core genomes. Species with large core genomes (> ∼3000 genes) are marked in red within the tree, highly diverse species (> ∼0.035) in blue. Tree labels and scatter plots are colored by their taxonomic annotations. Bottom panel: Relationships between habitats and core genome size and average nucleotide diversity of core genomes.

## Discussion

The question of how environments shape biological diversity is central to modern biology, extending beyond evolutionary biology. Microbial evolution is particularly affected by ecological constraints due to the broad distribution of microbial life across virtually all environments on Earth. Our understanding of microbial species and their evolution has been extended by the pan-genome concept [5, 6]. By analyzing microbial pan-genomes in the context of their environmental preferences and phylogeny, we were able to dissect major forces that shape microbial genomes. Our results suggest that habitat and phylogeny explain the majority of variation across different pan-genomic features, with differential contributions to size and divergence measures. Different theories and concepts have been postulated to explain microbial evolution in response to the environment. For example, it has long been thought that a large pool of accessory genes would be beneficial in certain habitats (and habitat combinations), but the role of the pan-genome as an adaptive evolutionary entity has been recently disputed. In the respective debate [8, 61–66], analyses of pan-genome size estimates (Fig. 2b) have led to the conclusion that pan-genomes are adaptive [8], while studies focusing on diversity measures such as genome fluidity led to the conclusion that pan-genome evolution is predominantly neutral [61]. Our analysis shows that environmental conditions and phylogenetic inertia affect size-related pan-genome features to a higher degree (than diversity features), suggesting that the adaptiveness of pan-genomes is at least partially explained by environmental preferences of species and their phylogenetic inertia (Fig. 2b). Mechanistically, it is likely that ecological constraints imposed by habitats drive pan-genome evolution, through natural selection, genetic drift and/or both and most likely in dependence on the species’ effective population size [67]. Yet, pan-genome size and other features are also partially determined by phylogenetic inertia: we observed that core genome size and average genome size (number of protein-coding genes) were most affected by phylogenetic position (Fig. 2b, Fig. 4). The conservation of the core genome in a given clade is likely due to the fact that it consists of essential genes that are under strong negative selection pressure [68–70], which leads to vertical “heritability” of its content and size from ancestral species to descendants during speciation events.

Building on a previous study, which showed a weak positive relationship between the ubiquity of species and overall genome size [58], we found that the strongest (albeit still moderate) correlation was with core genome size. Our more detailed observations suggest that genes that facilitate ubiquity (i.e., present across many habitats) are usually present in the core genome, which is further supported by the absence of a significant correlation between average intra-species pairwise functional distance and ubiquity (Fig. 2a). If functional diversity of accessory genome was highly important for ubiquity, we would expect a positive correlation between intra-species pairwise functional distance and ubiquity. In other words, the expansion of a species into additional habitats requires almost all strains to have genes that facilitate survival and proliferation in all or most species habitats.

Overall, our results indicate important relationships between the environment, macroevolutionary patterns, and microevolutionary (pan-genome) features, exemplified by the association between ubiquity and core-genome size. Hence, multi-feature predictive modelling is able to predict the ubiquity and environmental preferences of microbial species from pan-genomic information and phylogenetic placement, whereby accuracy will increase as more (pan)genomes become available. Functional knowledge of the genes within the pangenome will also help to predict habitat ranges as well as required or desired environmental conditions, in the context of the respective phylogenetic placements.

## Acknowledgements

The authors thank Bernd Klaus, Lucas Moitinho-Silva, Georg Zeller, Michael Kuhn, Thea van Rossum, Nassos Typas and Thomas Dandekar for advice with statistical analysis and discussion, Yan Yuan for maintenance of computational infrastructure. We thank João Matias Rodrigues for providing access and advice about Microbe Atlas Project database. O.M.M., F.H., T.S.B.S., S.S.L., P.B. were supported by European Research Council grant MicroBioS (ERC-2014-AdG). L.P.C. and J.H.-C. were supported by the European Union’s Horizon 2020 Research and Innovation Programme (grant #686070; DD-DeCaF). D.R.M. was supported by EMBO (ALTF 721-2015) and the European Commission (LTFCOFUND2013, GA-2013-609409). F.H. received funding through the European Union’s Horizon 2020 research and innovation program under the Marie Skłodowska-Curie grant agreement #660375.

