## Supplementary Figures for "Disentangling the impact of environmental and phylogenetic constraints on prokaryotic strain diversity"

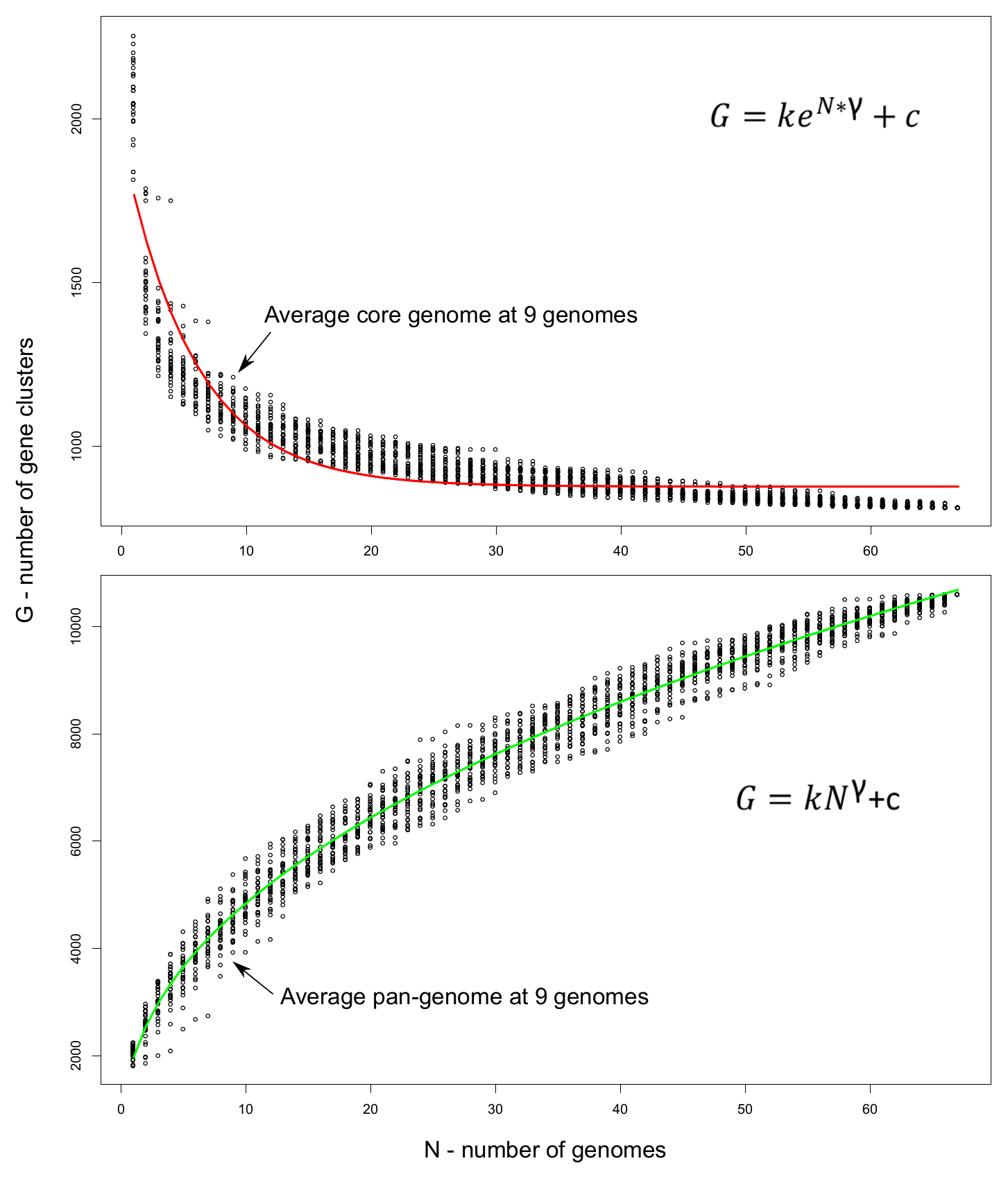

**Supplementary Figure 1.** Examples of saturation curves. Core- and pan-genome size at a given number of (randomly chosen) genomes from a species. Fits to displayed formulas shown in red (core genome) and green (pan-genome).

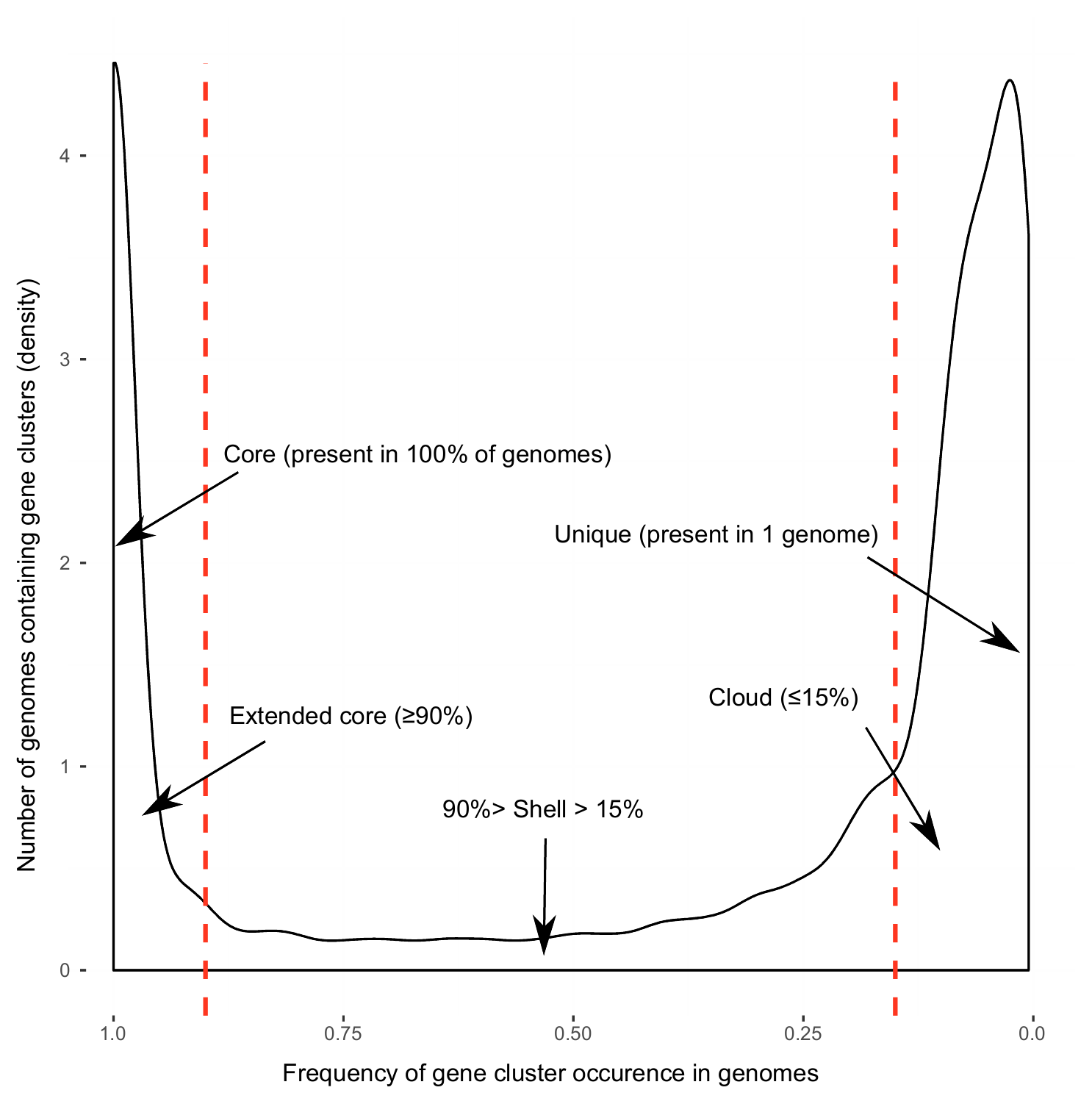

**Supplementary Figure 2.** Thresholds for pan-genome components. Gene frequency distribution displayed in black. Dash lines represent boundaries between extended core and shell genome; shell and cloud genome.

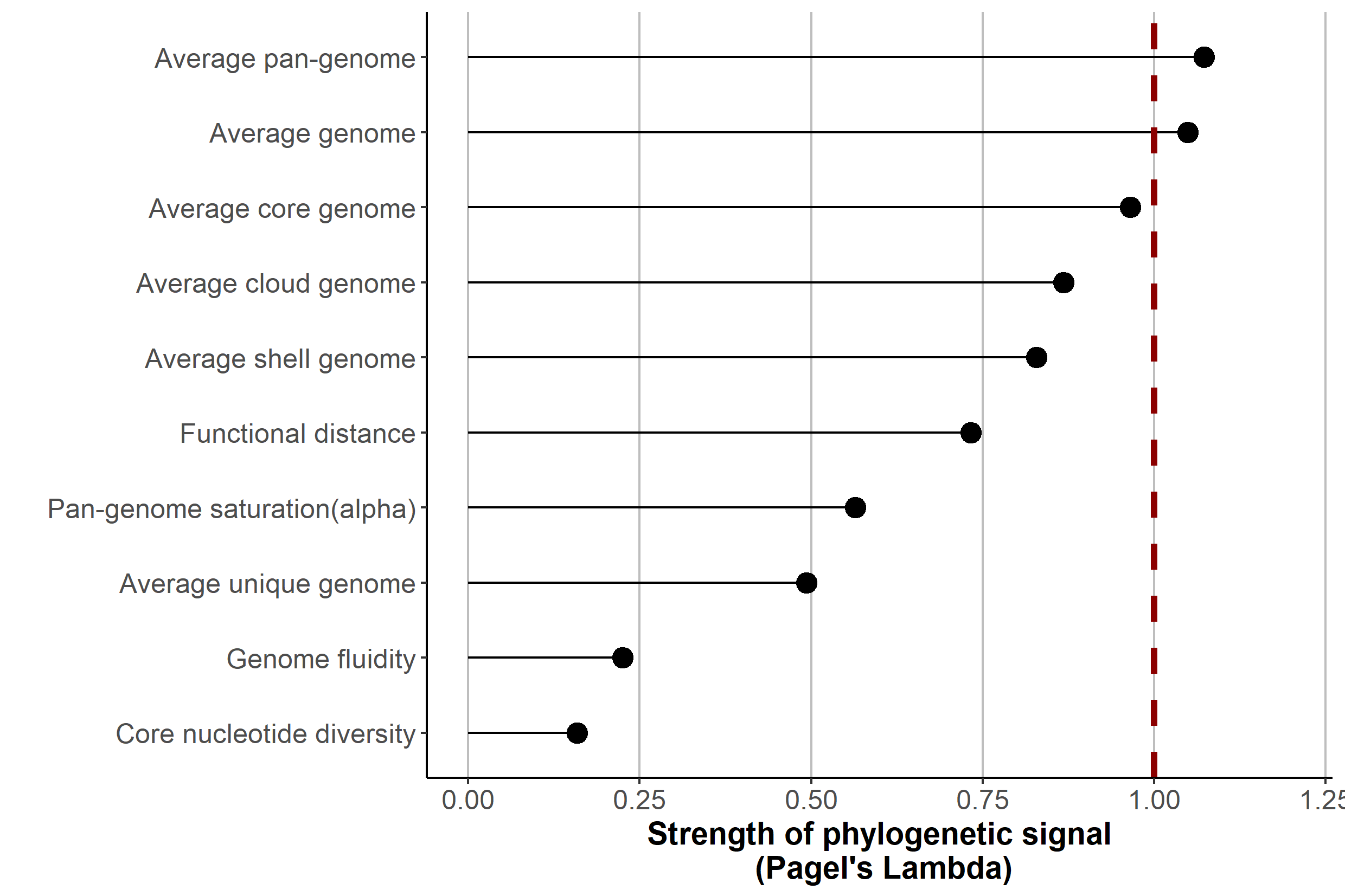

**Supplementary Figure 3**. Phylogenetic signal of 10 genomic characteristics across 155 species of Prokaryotes. When Pagel’s λ approximates to 1 – trait manifests phylogenetic signal (marked with dash line).

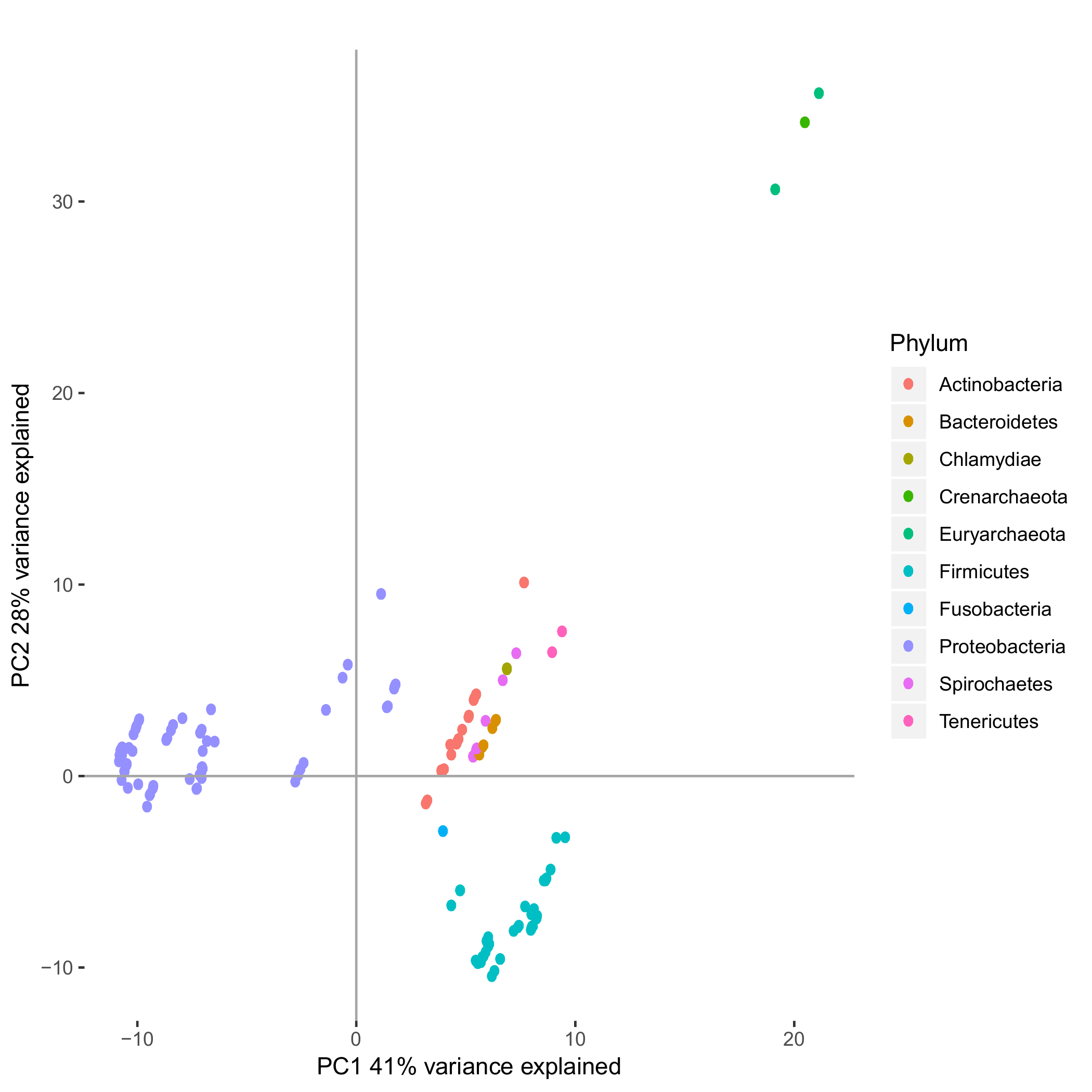

**Supplementary Figure 4.** Biplot of PCA (PC1 and PC2) using cophenetic distances observed in the phylogenetic tree reconstructed from 155 species used in this study.

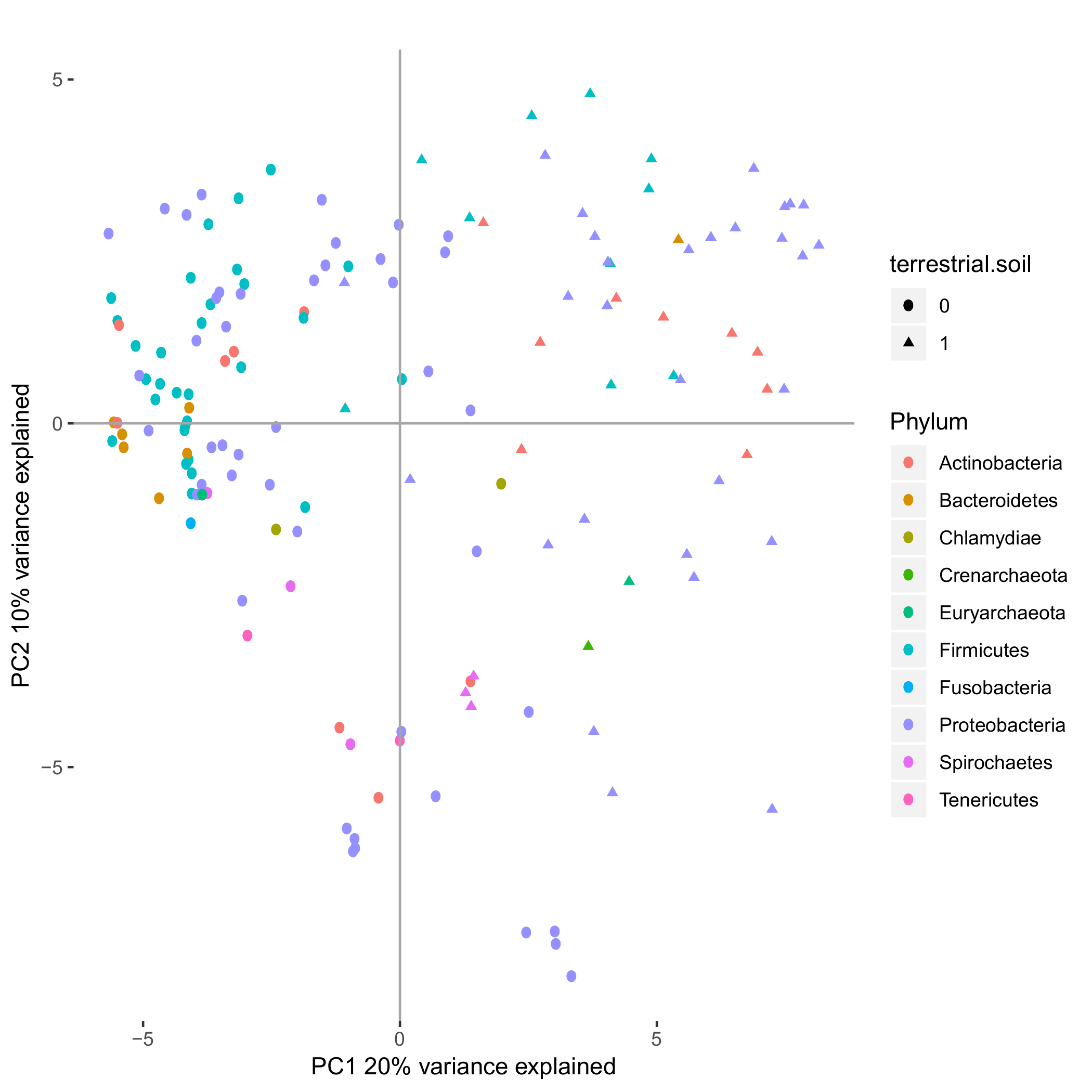

**Supplementary Figure 5.** Biplot of the first 2 principal components of the decomposition of the habitat-association matrix (0 - not associated with soil habitat, 1 - associated with soil habitat).

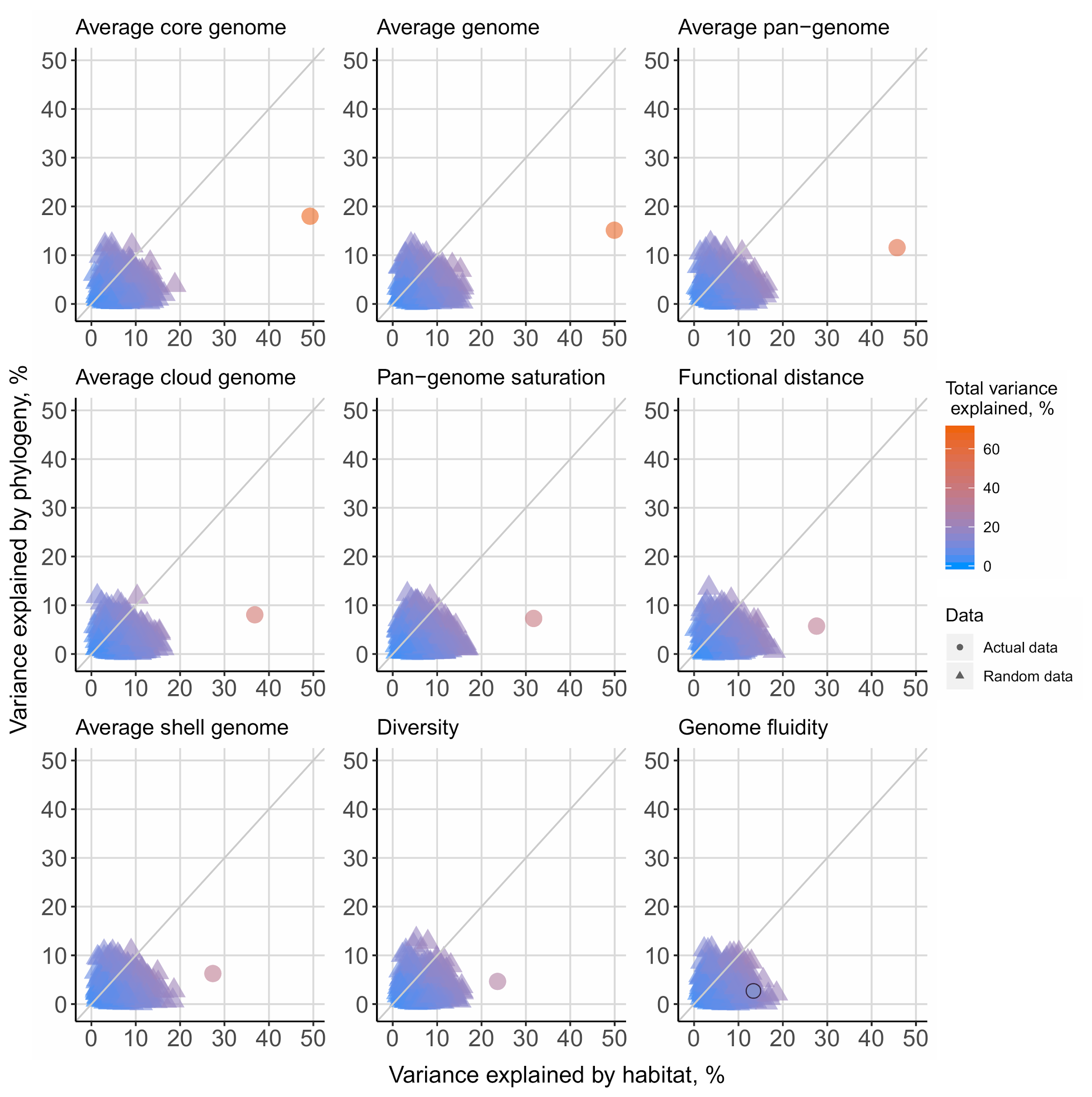

**Supplementary Figure 6**. Randomized phylogeny and habitat explain a smaller fraction of the variance than actual data for core genome size and average genome.
