## Supplementary Table 1 for "Disentangling the impact of environmental and phylogenetic constraints on prokaryotic strain diversity"

| Supplementary Table 1. Definitions of pan-genome features and other terms used in the manuscript | |
| --- | --- |
| Term | Definition |
| Number of genomes (sample size) | Number of genomes(also referred as sample size) that were used to calculate pan-genome features for each species |
| Pan-genome saturation(alpha) | Absolute value of alpha in equation [1] in methods |
| Pan-genome saturation(gamma) | Coefficient gamma from equation [2] in methods |
| Core genome saturation | Coefficient gamma from equation [3] in methods |
| Pan-genome | Total number of protein coding gene clusters estimated in entire number of genomes for each species |
| Core | Number of protein coding gene clusters in core genome calculated on entire number of genomes in each species |
| Extended core | Total number of protein coding gene clusters in extended core of pan-genome calculated on entire number of genomes for each species |
| Shell | Total number of protein coding gene clusters in shell genome of pan-genome calculated on entire number of genomes for each species |
| Cloud | Total number of protein coding gene clusters in cloud genome of pan-genome calculated on entire number of genomes for each species. Cloud genome also include unique gene clusters. |
| Unique | Total number of protein coding gene clusters in unique genome of pan-genome calculated on entire number of genomes for each species. |
| Average genome | Average number of protein coding gene clusters across all genomes in each species |
| Average extended core | Average number of protein coding gene clusters in extended core in an average genome of each species |
| Average shell | Average number of protein coding gene clusters in shell genome in an average genome of each species |
| Average cloud | Average number of protein coding gene clusters in cloud genome in an average genome of each species |
| Average unique | Average number of unique protein coding gene clusters in an average genome of each species |
| Average core genome | Average number of protein coding gene clusters in core genome in 9 genomes in each species |
| Average pan-genome | Average number of protein coding gene clusters in pan-genome on average in 9 genomes in each species |
| Pan-genome size (Chao lower bound estimate) | Chao lower bound estimate of possible number of genes in pan-genome function from "micropan" R-package) |
| Diversity | Average (1 - nucleotide identity) in core genomes of all genomes in each species |
| Average Jaccard distance | Average (1 - Jaccard index) of all genomes in each species |
| Genome fluidity | Genomic fluidity is the ratio of unique gene families to the sum of gene families in pairs of genomes averaged over randomly chosen genome pairs from within a group of *N* genomes |
| Functional distance | COG-based Average Jaccard distance between isolates/strains within species |
| Ubiquity of species | Ubiquity is the sum of all positive associations (Benjamini-Hochberg-corrected Fisher’s Exact Tests, p≤0.05) with each habitat in the Microbial Atlas Project dataset. In other words, ubiquity shows with how many habitats certain species was associated. |
